# Inferring tumor evolution from longitudinal samples

**DOI:** 10.1101/526814

**Authors:** Matthew A. Myers, Gryte Satas, Benjamin J. Raphael

**Affiliations:** Department of Computer Science, Princeton University, Princeton, NJ 08540.; Department of Computer Science, Brown University, Providence, RI 02912.

## Abstract

**Background:** Determining the clonal composition and somatic evolution of a tumor greatly aids in accurate prognosis and effective treatment for cancer. In order to understand how a tumor evolves over time and/or in response to treatment, multiple recent studies have performed longitudinal DNA sequencing of tumor samples from the same patient at several different time points. However, none of the existing algorithms that infer clonal composition and phylogeny using several bulk tumor samples from the same patient integrate the information that these samples were obtained from longitudinal observations.

**Results:** We introduce a model for a longitudinally-observed phylogeny and derive constraints that longitudinal samples impose on the reconstruction of a phylogeny from bulk samples. These constraints form the basis for a new algorithm, <u>C</u>ancer <u>A</u>nalysis of <u>L</u>ongitudinal <u>D</u>ata through <u>E</u>volutionary <u>R</u>econstruction (CALDER), which infers phylogenetic trees from longitudinal bulk DNA sequencing data. We show on simulated data that constraints from longitudinal sampling can substantially reduce ambiguity when deriving a phylogeny from multiple bulk tumor samples, each a mixture of tumor clones. On real data, where there is often considerable uncertainty in the clonal composition of a sample, longitudinal constraints yield more parsimonious phylogenies with fewer tumor clones per sample. We demonstrate that CALDER reconstructs more plausible phylogenies than existing methods on two longitudinal DNA sequencing datasets from chronic lymphocytic leukemia patients. These findings show the advantages of directly incorporating temporal information from longitudinal sampling into tumor evolution studies.

**Availability:** CALDER is available at https://github.com/raphael-group.

## 1 Introduction

Cancer is an evolutionary process, where cells accumulate somatic mutations over time through the process of clonal evolution [17]. As a result, many tumors contain heterogeneous populations of cells, or *clones*, with different combinations of somatic mutations. The clonal composition of a tumor may shift over time, particularly in response to treatment. Longitudinal sequencing, or sequencing DNA from tumor samples of the same patient at different time-points, is increasingly being used to track the progression of cancer over time. For example, Nadeu et al. [16] performed longitudinal DNA sequencing of 48 chronic lymphocytic leukemia (CLL) patients to study the impact of clonal and subclonal driver mutations on clonal dynamics and disease progression. In another study, Griffith et al [9] performed longitudinal sequencing of an acute myeloid leukemia (AML) patient and found a clone with a driver mutation in the gene IDH2 [9] that was present in less than 2% of the pre-treatment sample, but subsequently became the dominant clone after relapse. Longitudinal sequencing helps to identify and characterize clones that are present in small proportions prior to treatment, but become founder clones of a relapse or metastasis. Beyond blood cancers, longitudinal sequencing is increasingly being applied in clinical settings with the advent of noninvasive sampling of circulating tumor DNA (ctDNA) and circulating tumor cells (CTCs) – i.e., *liquid biopsies* [11]. Such studies allow researchers to track the evolutionary trajectories of cancer and hold promise for providing greater understanding of how tumor clones and individual mutations interact and respond to treatment.

A fundamental step in studying the evolutionary dynamics of cancer is the construction of a phylogenetic tree that describes the somatic mutation and cell division process. The vertices of such a tree correspond to cells (or clones) in the tumor, and edges describe ancestral relationships between cells/clones. The problem of constructing a phylogenetic tree from measurements of individual taxa is a well-studied problem. However, with DNA sequencing data of bulk tumor samples – each containing thousands to millions of cells – one must instead build a phylogenetic tree from *mixtures* of taxa. In recent years, a number of specialized algorithms have been developed to solve this phylogenetic mixture problem [1–5,12–14,18,19,22,24]. While these algorithms are designed to address the complexities of bulk sequencing data, none directly integrate the constraint that samples are measured longitudinally.

In this paper, we leverage the additional information provided by longitudinal measurements in the reconstruction of phylogenetic trees from bulk samples. First, we introduce a model for longitudinally-observed phylogenies as vertex-colored trees in which the order of colors encodes the temporal order of the samples. Next, we formulate the Longitudinal Variant Allele Frequency Factorization Problem (LVAFFP), a generalization of the problem of reconstructing a tree from bulk samples where the samples are temporally ordered. We derive a combinatorial characterization of the solutions to the LVAFFP as constrained spanning trees of a directed graph constructed from the variant allele frequencies (VAFs) of somatic mutations. We then introduce an algorithm, <u>C</u>ancer <u>A</u>nalysis of <u>L</u>ongitudinal <u>D</u>ata through <u>E</u>volutionary <u>R</u>econstruction (CALDER), to solve the LVAFFP both for perfect data and for data that has uncertainty in the observed frequencies. In data with uncertainty, we use confidence intervals to capture uncertainty in VAFs and a Lasso-like objective function to enforce sparsity on the proportions of clones present at each time point. We also describe how the longitudinal structure of the data is revealed by the presence and absence of clones and mutations, and introduce an absence-aware method for clustering mutations into clones.

We compare CALDER to several existing approaches [3, 13, 14] for tumor phylogeny reconstruction on simulated and real longitudinal sequencing data. On simulated data, we show that longitudinal constraints reduce the solution space of phylogenetic trees that describe the bulk sequencing data by up to 90%, and on average by 30%. We apply CALDER to longitudinal bulk DNA sequencing data from CLL patients from Rose-Zerilli et al. [20] and Schuh et al. [23]. We find that CALDER produces more plausible evolutionary trees that respect the longitudinal ordering of the data.

**Figure 1:**
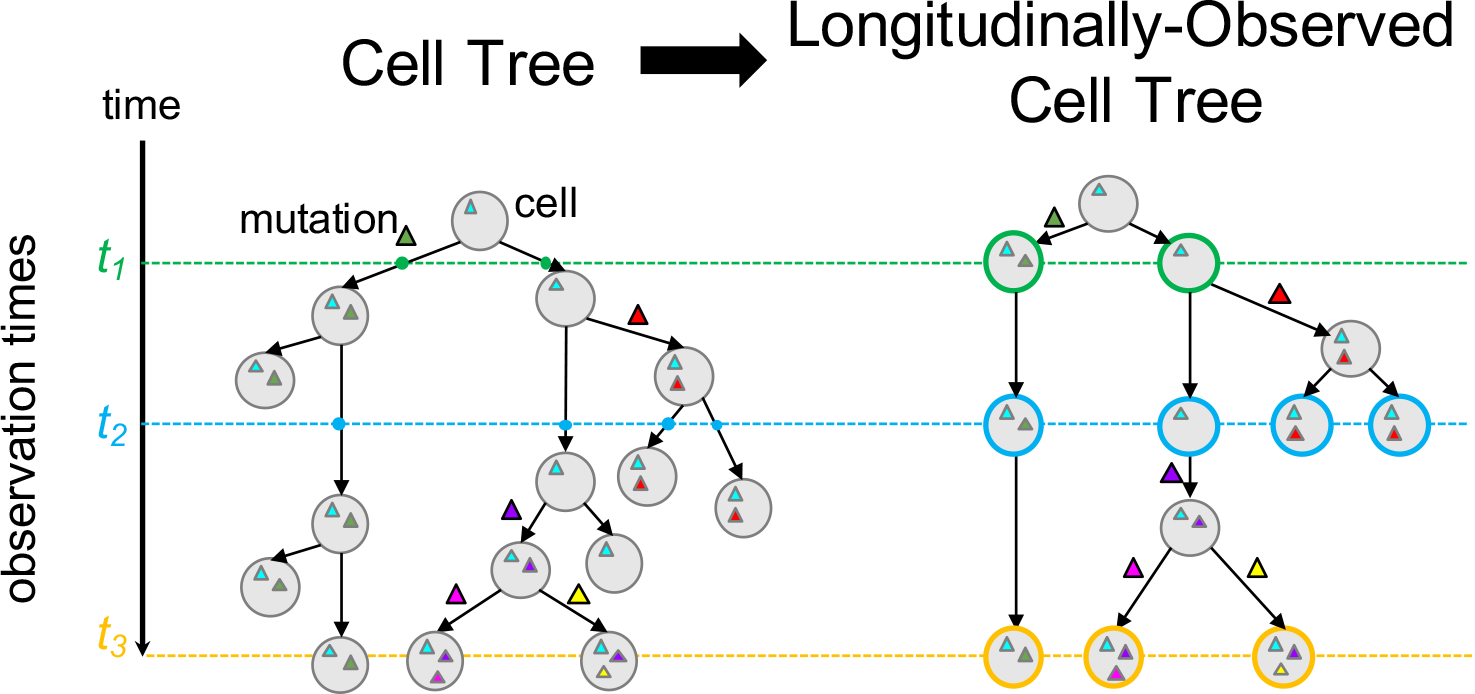
(Left) A *cell tree* describes the cell division and mutation history of a tumor, where vertices are cells and edges are parental relationships. We are typically not able to observe this evolutionary process continuously, and instead have a set of observations from discrete times. (Right) A *longitudinally-observed cell tree* describes the ancestral relationships between the observed cells. Cells are labeled (colored) with the time that they are observed.

## 2 Methods

### 2.1 Longitudinal observation of cancer evolution

The evolutionary history of a tumor at the level of individual cells can be represented by an edge-weighted binary *cell tree P* = (*V, E*), where a vertex *v* ∈ *V* is a cell, a directed edge between two cells indicates a parental relationship, the weight of an edge indicates the lifespan of a cell, and edges are labeled by the mutation(s) that distinguish child from parent. In longitudinal data, we observe a cell tree at a set of discrete time-points *t*_1_ < *t*_2_ < … < *t_m_* during the evolutionary process. Let *C*_1_, …, *C_m_* be the disjoint sets of cells present at times *t*_1_*, … t_m_* respectively. We define a *longitudinally-observed cell tree P* = (*V, E, c*) to be a vertex-colored tree where:

1. Every vertex *v* ∈ *V* has a color *c*(*v*) ∈ {0, …, *m*} where *c*(*v*) = *i* indicates that *v* ∈ *C_i_*, and *c*(*v*) = 0 indicates that *v* is not observed;
2. For every path *p* = *r*, …, *v* from the root *r* to a leaf *v*, the nonzero colors on the path are encountered in numerical order.

With bulk sequencing data, we do not observe individual cells directly and instead reconstruct tumor evolution at the level of *clones*, or populations of cells that share a set of mutations. Thus, instead of a cell tree, we build a *longitudinally-observed clone tree P* = (*V, E, c*) where each vertex *v* ∈ *V* corresponds to a distinct population of cells, edges (*v, w*) ∈ *E* are ancestral relationships, and colors *c* follow the properties stated above.

### 2.2 Reconstructing longitudinal phylogenies

Suppose we obtain DNA sequencing data from bulk tumor samples at times *t*_1_ < *t*_2_ < … *t_m_*. Across the samples, we identify a total of *n* genomic positions containing single-nucleotide somatic mutations. For each sample *t* and each mutation *i*, we compute the *variant allele frequency (VAF) f_t,i_* as the fraction of reads from sample *t* that cover position *i* and contain mutation *i*. In the absence of copy-number aberrations, the variant allele frequency *f_t,i_* is directly proportional to the proportion of cells in sample *t* that contain mutation *i*. We represent the observed data as a *frequency matrix F* = [*f_t,i_*]. Following El-Kebir et al. [3], we assume that each mutation corresponds to a clone (i.e., we assume that mutations have been clustered according to their VAFs [13, 15, 21]). Each tumor sample *t* is then a mixture of clones, where *u_t,i_* is the mixing proportion of clone *i* in sample *t*. Thus, *u_t,i_* ≥ 0 and 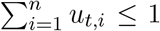. We define the *m* × *n* usage matrix *U* = [*u_t,i_*].

The fundamental problem in reconstructing a phylogenetic tree from bulk tumor samples is to identify the clone tree *T* and usage matrix *U* that generated the observed mutation frequencies *F*. Multiple algorithms have been developed to solve this problem in various settings [1–5, 12–14, 18, 19, 22]. All of these algorithms rely on the infinite sites assumption (ISA), which states that a mutation is gained at most once during evolution and is never subsequently lost. Under the ISA – also known as the perfect phylogeny model – there is a 1-1 correspondence between a phylogenetic tree *T* and a 01-valued matrix *B*. Using this 1-1 correspondence, El-Kebir et al. [3] formulated the problem of reconstructing a phylogenetic tree from bulk tumor samples as a matrix factorization problem.

#### Variant Allele Frequency Factorization Problem (VAFFP)

*Given frequency matrix F, find perfect phylogeny matrix B and usage matrix U such that F* = *UB*.

The *n* × *n* matrix *B* is a binary perfect phylogeny matrix [10] whose rows correspond to clones and columns correspond to mutations, where *b_i,j_* indicates if clone *i* has mutation *j*. Under the ISA, there is a 1-1 correspondence between perfect phylogeny matrices *B* and *mutation trees T* = (*V_T_, E_T_*) which describe the evolutionary relationships between clones. Vertices *v* ∈ *V_T_* correspond to clones, and edges (*v, w*) ∈ *E_T_* correspond to parental relationships between clones. We label each edge of a mutation tree with the mutation(s) that distinguish the parent and child vertex. Let *M_v_* be the set of mutations present in the clone corresponding to vertex *v*. Edge (*v, w*) ∈ *E_T_* is labeled by the difference *M_v_*\ *M_w_* in mutations between the clones. Under the ISA, each mutation appears at most once on an edge in *T*. Let *μ* be a mapping from each vertex *v* ∈ *V_T_* to the mutation on its incoming edge, i.e., *μ*(*v*) = *j*. Without loss of generality, we use a single phylogenetic character to represent the set of mutations on each edge. In contrast to a clone tree, a mutation tree has exactly one vertex corresponding to each clone. Mutation trees are analogous to *n*-clonal trees as described in [3]. Under the ISA, the clones that contain the mutation *j* are the first clone *v* that acquired mutation *j* (i.e., *μ*(*v*) = *j*), and the descendants Δ*_v_* of *v*. Thus, we have that the frequency 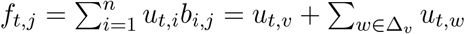.

The usage matrix *U* and perfect phylogeny matrix *B* (together with *B*’s corresponding mutation tree *T*) determine a vertex-colored *observed clone tree P_U,B_*, which we construct as follows. See Figure 2 for an example.

1. **Add observed clones.** For each vertex *v* ∈ *T* and the corresponding column *j* = *μ*(*v*) of *U*, we represent the observations of *v* in the observed clone tree as follows. Let *t*_*i*_1__ < *t*_*i*_2__ < … < *t*_*i*_*k*__ be the nonzero entries in column *j* of *U*. Add path *π*(*v*) = *v*_0_ → *v*_*i*_1__ → … → *v*_*i*_*k*__ to *P_U,B_*, such that *c*(*v*_0_) = 0, and *c*(*v*_*i*_*l*__) = *t*_*i*_*l*__ for *l* = 1 … *k*.
2. **Add mutation edges.** Let 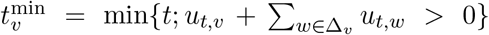, where Δ*_v_* is the set of descendants of *v* in *T*. For each edge (*v,w*) in *T*, add edge (*v′,w′*) to *P_U,B_*, where *v′* = 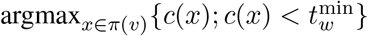 and *w′* is the uncolored vertex corresponding to clone *w*.
3. **Remove extra uncolored vertices.** For every uncolored vertex *v* with incoming edge (*u, v*) labeled with mutation *i*, and exactly one *unlabeled* outgoing edge (*v, w*), replace vertex *v* and edges (*u, v*) and (*v, w*) with a single edge (*u, w*) labeled with mutation *i*.

**Figure 2:**
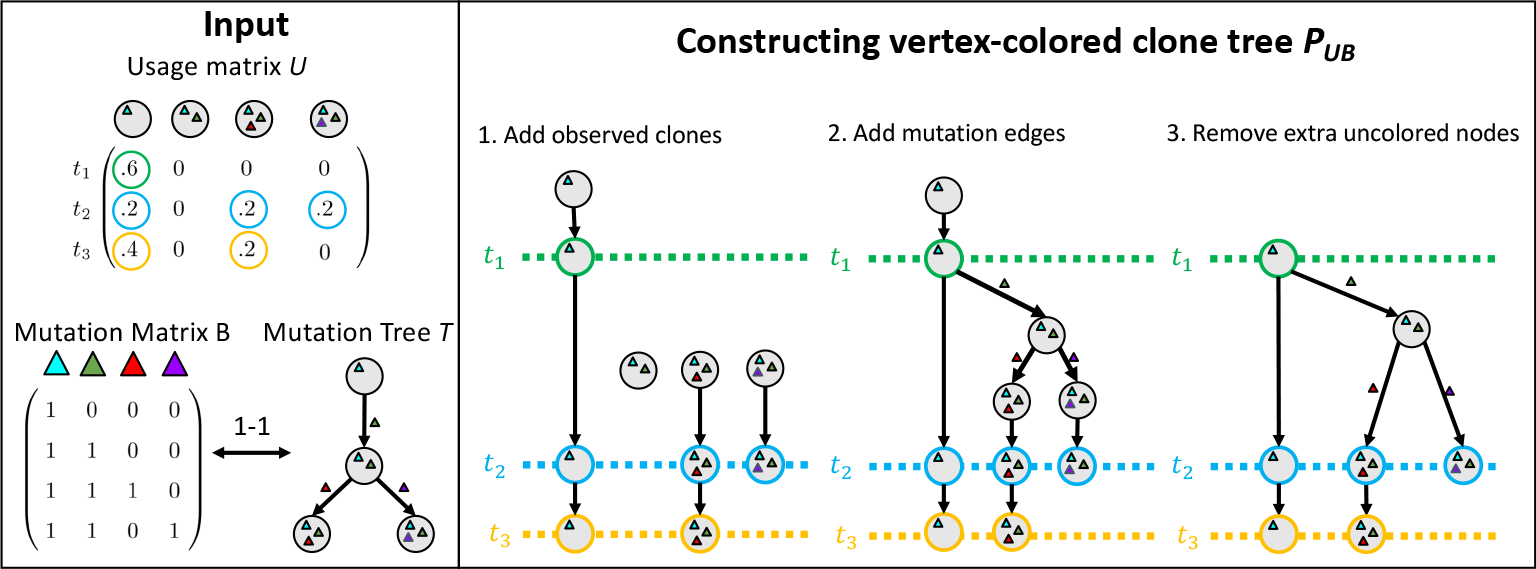
Construction of the observed clone tree *P*_*U*,*B*_ from usage matrix *U* and perfect phylogeny matrix *B*.

The colored vertices in *P_U,B_* correspond to nonzero entries in *U*, and the edges in *P_U,B_* maintain the ancestral relationships in *T*: if mutation *i* precedes mutation *j* in *T*, then *i* also precedes *j* in *P*. In general, there are multiple clone trees that could be generated from a *U* and *B* – for example, one could add arbitrarily many unobserved clones with corresponding uncolored vertices. However, the construction above gives a unique clone tree *P_U,B_*. Moreover, if there exists a longitudinally-observed clone tree corresponding to the factorization (*U, B*), then the *P_U,B_* constructed above is longitudinally observed.

In the VAFFP, there are no assumptions regarding the temporal relationships between bulk sequencing samples. Indeed, often there are multiple factorizations *F* = *UB* for a given frequency matrix *F*. Interestingly, given a frequency matrix *F* obtained from longitudinal sampling, it is possible to obtain a factorization *F* = *UB* where the observed clone tree *P_U,B_* is not longitudinally observed (Figure 3). Thus, given DNA sequencing data from longitudinal bulk tumor samples, the problem one wants to solve is the following variation of the VAFFP.

**Figure 3:**
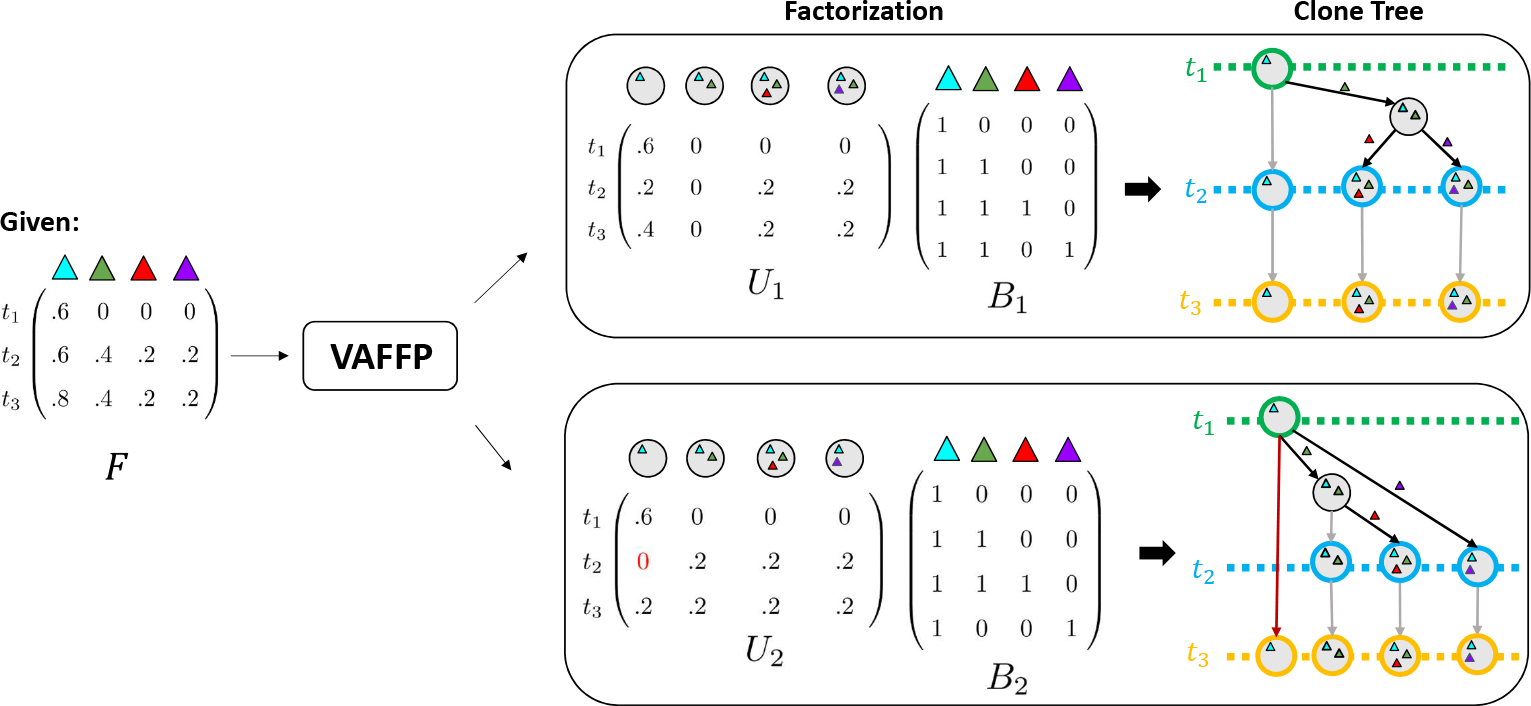
Not all VAFFP solutions correspond to longitudinally-observed clone trees. An example of a frequency matrix *F* which yields two factorizations. The factorization *F* = *U*_1_*B*_1_ (top) corresponds to a longitudinally-observed clone tree, but the factorization *F* = *U*_2_*B*_2_ (bottom) does not. This is evident in the red highlighted path, which does not respect the ordered coloring.

#### Longitudinal Variant Allele Frequency Factorization Problem (LVAFFP)

*Given frequency matrix F whose rows correspond to longitudinal samples, find a perfect phylogeny matrix B and usage matrix U such that F* = *UB and the observed clone tree P_U,B_ is longitudinally observed*.

This problem, like the VAFFP, is NP-complete (see proof in Supplemental Information section 1).

### 2.3 Characterizing solutions to the LVAFFP

For a frequency matrix *F*, let 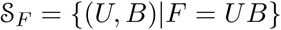 be the set of solutions to the VAFFP, and let 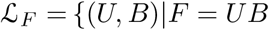 and *P_U,B_* is longitudinally observed be the set of solutions to the LVAFFP. Note that for all frequency matrices *F*, 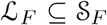. El-Kebir et al. [3] and Popic et al. [18] have previously characterized 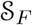 as a set of constrained spanning trees on a particular directed graph, and describe algorithms to enumerate these trees. Thus, one approach to solving the LVAFFP is to enumerate all trees corresponding to solutions in 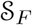 (solutions to the VAFFP) and check whether the corresponding clone trees are longitudinally observed. In the following, we describe a procedure to enumerate 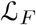 directly.

First, we review the necessary and sufficient conditions that characterize solutions 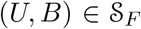; these conditions were presented in [3, 18]. For a vertex *v* ∈ *T*, let *δ_v_* be the children of *v*. In order for *T* to determine clones with non-negative mixture proportions, the following condition must hold for all samples *t* and mutations *v*,

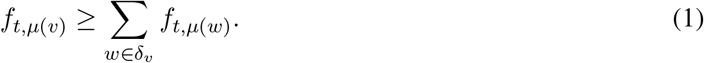

This is known as the Sum Condition in [3]. A frequency matrix *F* and mutation tree *T* = (*V, E*) (corresponding to perfect phylogeny matrix *B*) satisfying the Sum Condition *uniquely* determine a usage matrix *U* with *F* = *UB*, as follows:

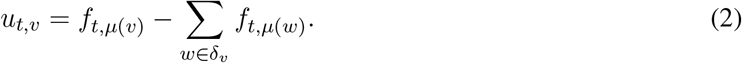

We will say that a tree *T* is a solution to the VAFFP (i.e., 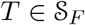) provided that the *U* defined in Equation 2 has non-negative entries.

Figure 3 shows that there are frequency matrices *F* for which 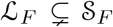. Thus, we are interested in deriving conditions on *F* that characterize 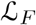. First, we derive conditions on a mutation tree *T* and usage matrix *U* for the corresponding clone tree *P_U,B_* to be longitudinally observed. For a mutation tree *T*, let Δ*_v_* be the set of vertices in the subtree of *T* rooted at *v*. For a clone *v*, let 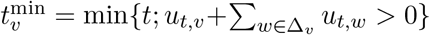 represent the first sample after its birth, and let 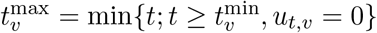 represent the first sample after its death. We have the following result.

#### Lemma 1.

*The following conditions are necessary and sufficient for a usage matrix U and mutation tree T to determine a longitudinally-observed clone tree P:*

1. ***Permanent extinction****. For all clones v, u_t,v_* = 0 *for all* 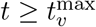.
2. ***Lineage continuity***. *For each edge* 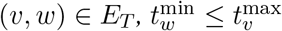.

Using the relationship between usage matrix *U* and frequency matrix *F* (Equation 2), we can express 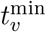 and 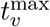 directly in terms of the frequency matrix as follows:

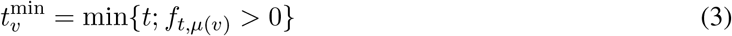

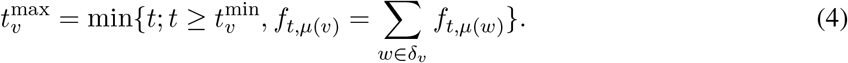

Note that Δ*_v_* disappears from the definition of 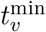, as mutation *μ*(*v*) is shared by all clones *w* ∈ Δ*_v_*, and that the strict equality in the definition of 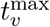 directly corresponds to a mixture proportion of 0.

Then, the necessary and sufficient conditions on *F* for a mutation tree *T* = (*V, E*) and a usage matrix *U* to determine a longitudinally-observed clone tree are as follows:

1. **Sum condition (SC)**. For all clones *v* and samples *t*, 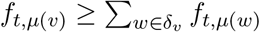.
2. **Permanent extinction condition (PEC)**. For all clones *v*, 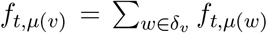 for all samples 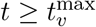.
3. **Lineage continuity condition (LCC)**. For each edge 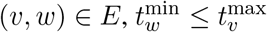.

### 2.4 Trivially longitudinal factorizations

It is not difficult to see that only the zero entries in *U* determine whether a pair *B, U* correspond to a longitudinally-observed clone tree, as stated in the following Lemma.

#### Lemma 2.

*If F* = *UB for some perfect phylogeny matrix B and usage matrix U, and U is strictly positive (i.e., u_t,p_* > 0 *for all time points t and clones p), then the observed clone tree P_U,B_ is longitudinally observed*.

As a result, we refer to solutions of the LVAFFP with strictly positive *U* as *trivially longitudinal*. See Supplemental Information Figure 1 for an example. Such trivially longitudinal solutions are especially problematic when there is uncertainty in *F* (see below).

### 2.5 Enumerating spanning trees

In this section, we use the three constraints from the previous section – i.e., the sum condition, permanent extinction condition and lineage continuity condition – to enumerate the set 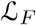 of solutions to the LVAFFP. Following [3, 18], we define the *ancestry graph G_F_* = (*V, E*) for a frequency matrix *F* to be the directed graph with vertices *V* = {1, …, *n*} and edges *E* = {(*p, q*)|*f_t,p_* ≥ *f_t,q_* for all *t* ∈ [1, *m*]}. The set 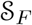 of solutions to the VAFFP correspond to the set of spanning trees of *G* where the sum condition (SC) is met at each vertex [3, 18]. Similarly, the set 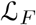 of solutions to the LVAFFP is the set of spanning trees where the sum condition (SC), permanent extinction condition (PEC) and lineage continuity condition (LCC) are met.

We enumerate the set 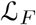 by adapting the Gabow-Myers algorithm [7] as was done previously for the VAFFP and extensions [4, 18, 22]. The Gabow-Myers algorithm iteratively builds a spanning tree by considering the addition of a single edge to the growing tree using depth-first exploration and backtracking. In our adapted algorithm, we add an edge to the tree only if for the resulting subtree *T′*, the SC, PEC and LCC are met. See Supplemental Information section 4 for the full details of the algorithm. This adaptation relies on the property of *rooted subtree consistency* for these conditions.

#### Definition 1

(Rooted Subtree Consistency). *A condition, a function that maps a tree to a truth value, is* rooted subtree consistent *provided that if a rooted tree T meets the condition, any subtree of T with the same root also meets the condition*.

Specifically, this property guarantees that we can enumerate the full set of constrained spanning trees by adding one edge at a time. We show that the SC, PEC, and LCC are rooted subtree consistent (see Supplemental Information).

### 2.6 Solving the LVAFFP on real data

In the previous sections, we assumed that the observed mutation frequencies *F* were the true mutation frequencies. On real data, however, there are two complications: (1) uncertainty in the variant allele frequencies (VAFs) estimated from aligned sequence reads, and (2) incomplete sampling. Tumor phylogenetic algorithms address the first issue using generative probabilistic models of observed read counts [2, 13, 22] or confidence intervals [3, 4, 14, 18, 24]. The second issue occurs because the clones that are measured in the sample may not be all the clones that are present at that time-point; for example, clones present in low proportions or in different spatial locations may not be quantified. Because our analysis relies on the presence and absence of clones in samples – corresponding to nonzero and zero values in *U*, respectively – failure to measure a clone may lead to incorrect phylogenies.

In order to accommodate uncertainty in the values in *F*, we derive a confidence interval for each entry *f_t,i_* from the number of reads covering position *i*. Let *F*^−^ be the matrix of lower bounds on frequency values, and let *F*^+^ be the matrix of upper bounds. Given *F*^−^ and *F*^+^, our goal is to infer a frequency matrix 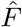, usage matrix *U*, and perfect phylogeny matrix *B* such that 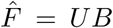, where *P_U,B_* is longitudinally observed and 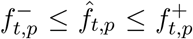 for all time-points *t* and samples *p*. With confidence intervals on mutation frequencies, longitudinal constraints become easy to satisfy. Specifically, for a fixed mutation tree *T*, the usage value *u_t,p_* for a clone *p* at time-point *t* satisfies 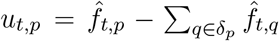, and thus is equal to the slack between the frequency of the mutation on the incoming edge and the sum of the frequencies of the mutations on the outgoing edges. When frequencies are adjustable within their confidence intervals, it is almost always possible to set 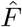 so that most or all usage values are strictly positive with very small values. These trivially longitudinal solutions are generally uninteresting, as they imply that all clones are present at all time-points even if the inferred proportions of these clones are miniscule.

To avoid such trivial solutions, we impose two additional constraints. First, we aim to minimize the number of non-zero entries in *U*. We do this by adding a regularization term |*U*|_1_ to the objective function. This term encourages sparsity in *U*, as in Lasso regression [25]. Second, we require that all non-zero entries of *U* exceed a minimum threshold *h*; that is, we require that *u_t,p_* = 0 or *u_t,p_* ≥ *h* for all *t, p*. These extra constraints prevent solutions with trivially small usages or solutions where the absence of a clone would violate the longitudinal constraints. Incorporating these extra constraints, we formulate the following problem.

#### Interval Longitudinal Variant Allele Frequency Factorization Problem (I-LVAFFP)

*Given frequency matrices F*^−^ *and F*^+^, *find a frequency matrix* 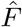, *usage matrix U, and perfect phylogeny matrix B such that:* 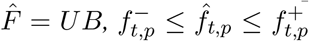 *and u_t,p_* ∉ (0, *h*) *for all time-points t and samples p*, |*U*|_1_ *is as small as possible, and P_U,B_ is longitudinally observed*.

We solve this problem using an ILP (Supplemental Information section 5).

### 2.7 Absence-Aware Clustering

When there is uncertainty in variant allele frequencies, there are typically sets of mutations whose frequencies are indistinguishable. The standard approach to address this issue is to cluster mutations into clones either prior to [15, 21, 26] or simultaneously with phylogeny reconstruction [2, 12, 13, 22]. In real data, however, it may difficult to distinguish between mutations that are present in low proportions and mutations that are absent from a sample. Clustering algorithms do not explicitly distinguish between these two classes, and thus may group mutations that are present in low proportion in a sample into the same cluster as mutations that are absent from a sample. Indeed, we have observed this behavior on real data using popular mutation clustering algorithms [15, 21]. Such clustering errors are particularly problematic in the analysis of longitudinal data, where the presence and absence of clones in different samples determine whether a specific phylogenetic tree is consistent with longitudinal samples.

We develop an *absence-aware* approach to cluster mutations into clones. Specifically, this approach relies on the principle that if a pair of mutations is present in the same set of clones, then they will be present in the same set of samples. Thus, prior to clustering, we partition the set of mutations into subsets where each subset of mutations is present in the same set of samples. Then we perform clustering independently on each of these subsets of mutations. Further details are in Supplemental Information section 6.

## 3 Results

### 3.1 Longitudinal constraints reduce ambiguity on simulated data

First, we use simulated data to assess what fraction of solutions to the VAFFP are longitudinal, i.e., solutions to the LVAFFP. We generate 10 mutation trees using the simulation procedure described in El-Kebir et al. [5], with between 4 and 13 vertices each as determined by the tumor evolution. For each of these trees, we generate frequency matrices *F*, for *m* = 2, …, 9 samples; this results in 10^6^ *F* matrices for each tree and value of *m*, and a total of 8 × 10^7^ *F* matrices (further details are in Supplemental Information section 7). We solve the VAFFP and the LVAFFP for each *F*, counting the number 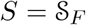 of solutions to the VAFFP and the number 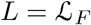 of solutions to the LVAFFP. Note that because all solutions to the LVAFFP are also solutions to the VAFFP, *L* ≤ *S*. Because we are primarily interested in the difference between *L* and *S*, we restrict our analysis to those instances for which *L* < *S*. We find that on the 423,328 instances with *L* < *S* (representing 0.5%-11% of the instances for each tree; mean 5.3%, median 3.9%), *L* ≈ 0.7*S* on average (and also median), with a minimum of *L* ≈ 0.07*S* (Figure 4).

**Figure 4:**
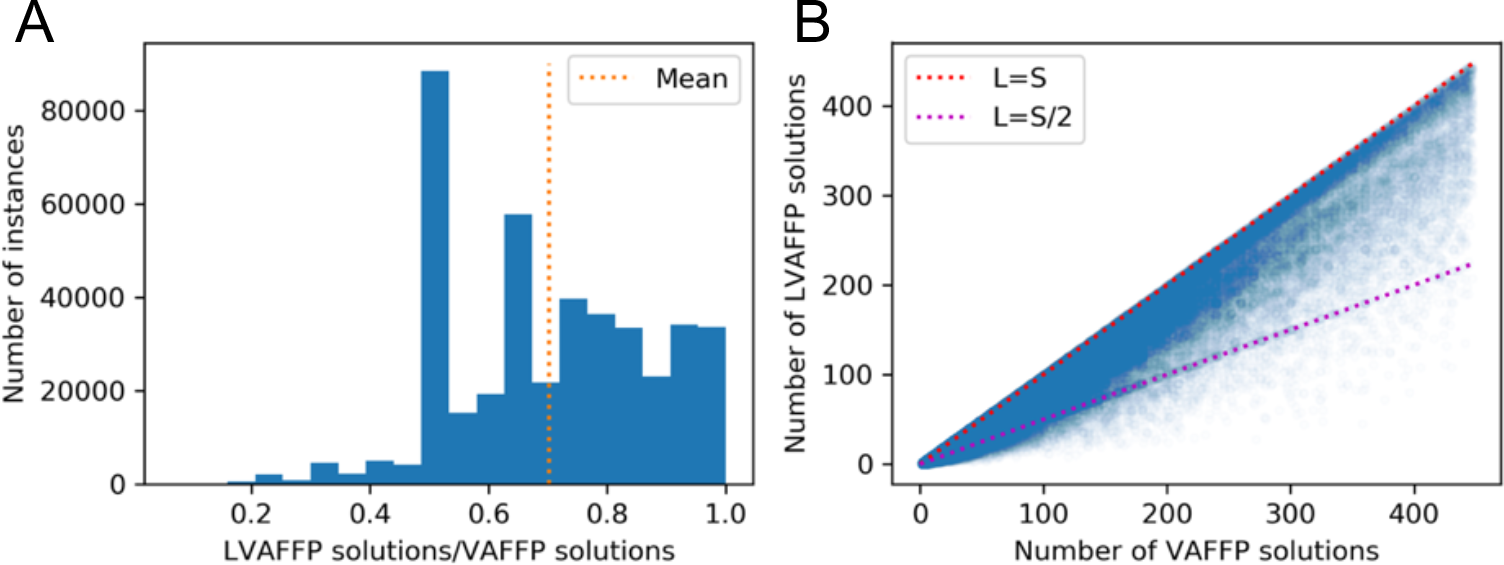
Comparison of the number *S* of solutions to the VAFFP and the number *L* of solutions to the LVAFFP on 423,328 simulated *m* × *n* frequency matrices which were generated uniformly at random from a total of 10 tree topologies, with *m* = 2, …, 9 and *n* = 4, …, 13. (A) Histogram showing the proportion of solutions to the VAFFP which are also solutions to the LVAFFP. (B) Scatterplot showing how the relationship between *S* and *L*. Note that we exclude instances where *L* = *S*.

### 3.2 CALDER analysis of longitudinal CLL sequencing data

We apply CALDER to longitudinal sequencing data from 13 chronic lymphocytic leukemia (CLL) patients from Rose-Zerilli et al. [20]. We find that longitudinal constraints result in fewer phylogenetic trees (i.e., *L* < *S*) for 4 of the 7 patients that were sampled at more than 2 time-points. Figure 5 shows CALDER results on patient 9, whose tumor was sequenced at 4 time-points during treatment, using targeted deep sequencing of 21 mutations that were identified at diagnosis. We excluded 6 mutations with VAF ≫ 0.5 from analysis as these likely overlap copy-number aberrations – further details on data processing are in Supplemental Information section 9. CALDER finds 110 maximal mutation trees with 9 mutations, 74 of which yielded longitudinally-observed clone trees. Figure 5B shows an example mutation tree *T*_1_ and the corresponding longitudinally-observed clone tree *P*_1_.

**Figure 5:**
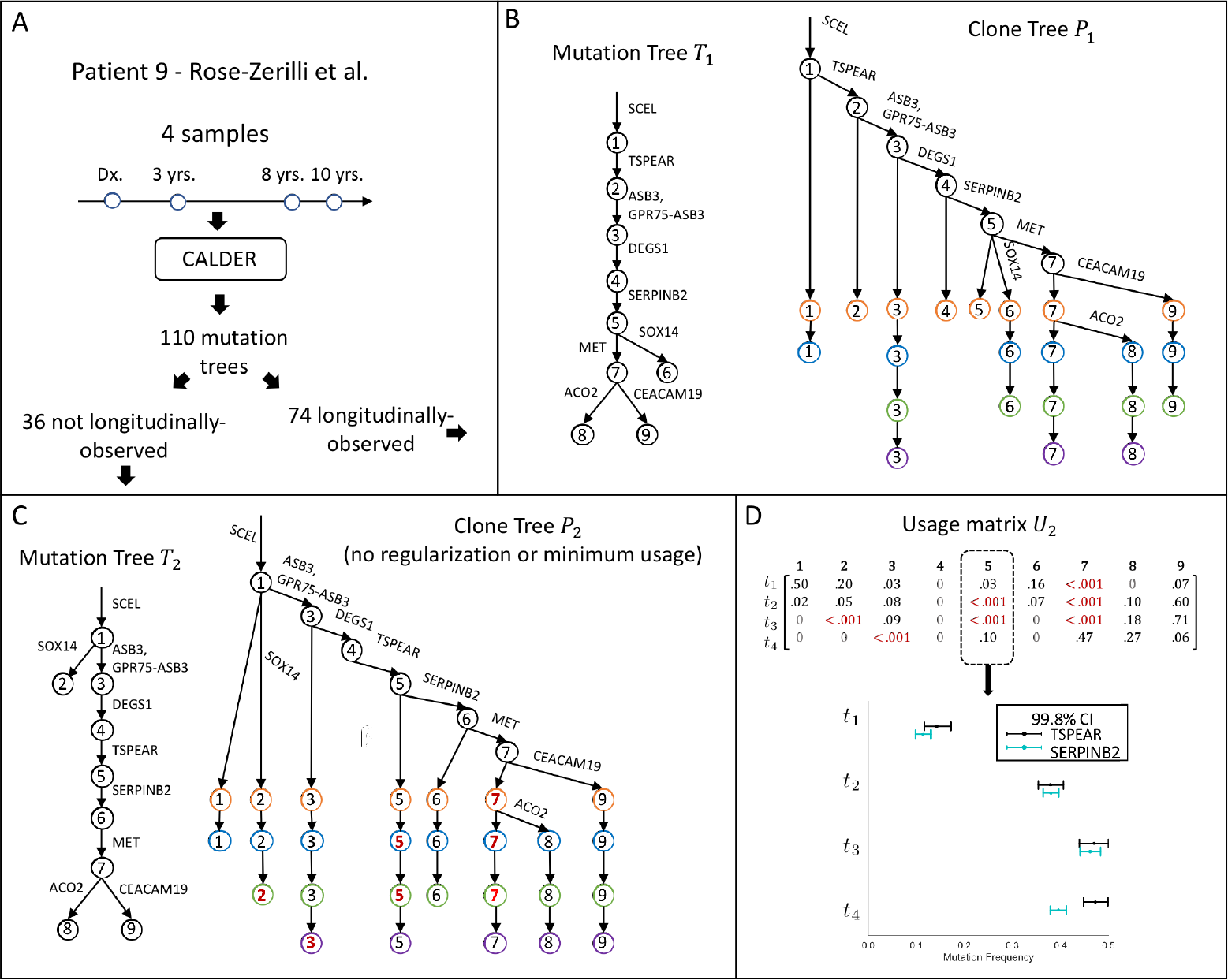
Longitudinal constraints reduce ambiguity on real data. CALDER results on CLL patient 9 from Rose-Zerilli et al. [20]. (A) 110 mutation trees are consistent with the mutation frequencies measured in this patient, but only 74 of these trees correspond to longitudinally-observed clone trees. (B) Example of a mutation tree *T*_1_ and its corresponding longitudinally-observed clone tree *P*_1_. (C) A mutation tree *T*_2_ that does not correspond to a longitudinally-observed clone tree. Without the requirement of a minimum clone usage *h*, we obtain the longitudinally-observed clone tree *P*_2_. (D) Usage matrix *U*_2_ corresponding to *T*_2_ has many small entries *u*_*ij*_ < 0.001 (highlighted in red). These small usages are required to meet longitudinal constraints. The support for the presence of clone 5 at each time-point *t*_*i*_ is the difference in frequency between mutations TSPEAR and SERPINB2. With the exception of *t*_4_, this difference is within the confidence bounds of each mutation, and thus clone 5 is unlikely to be present at times *t*_1_, *t*_2_, and *t*_3_.

To demonstrate the importance of the regularization term |*U*| _1_ and the minimum clone usage *h* in the I-LVAFFP, we examine in more detail a mutation tree *T*_2_ for which CALDER did not report a longitudinally-observed clone tree. We compute a usage matrix *U*_2_ and a mutation tree *T*_2_ that minimize 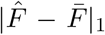, where the entries of *F̄* are means of the mutation frequency confidence intervals, such that 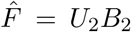 and the corresponding clone tree *P*_2_ is longitudinally observed. Figure 5C shows the mutation tree *T*_2_ and the corresponding longitudinally-observed clone tree found by this approach. This resulting usage matrix *U*_2_ has many entries *u*_*ij*_ < 0.001 (Figure 5). In particular, clone 5 is inferred to be present at all time points in order to meet longitudinal constraints, but has small usages (< 0.001) at time points *t*_2_ and *t*_3_ (Figure 5D). From the tree *T*_2_, the usage of clone 5 in a sample is equal to the difference in frequencies between mutations TSPEAR and SERPINB2. While the frequencies of these mutations are well separated at *t*_4_, they are indistinguishable (within the confidence intervals) at *t*_2_ and *t*_3_. Thus, the sequencing data does not support the presence of these clones, implying that the tree *T*_2_ is implausible. This example demonstrates how CALDER leverages longitudinal constraints to reduce ambiguity in clonal composition and clone tree reconstruction.

### 3.3 CALDER infers more plausible trees on longitudinal data than existing methods

We apply CALDER to a CLL patient CLL003 from Schuh et al. [23] that was previously analyzed in the papers introducing the PhyloSub [13], CITUP [14], and AncesTree [3] tumor phylogeny algorithms. For this patient, high-depth targeted sequencing was performed at five time-points, and the published analysis reported 20 mutations. We excluded 3 mutations whose VAFs were ≫ 0.5 as these likely overlap with copy-number aberrations – note that the published results from AncesTree [3] also exclude the same 3 mutations. We cluster mutations as described in Section 2.7, obtaining 4 mutation clusters which we label A, B, C, and D. CALDER infers a clone tree containing 4/5 clusters and 15/17 mutations (Figure 6A). This clone tree is longitudinally observed and shows a process of ongoing clonal evolution in this patient, with mutations in cluster B occurring between *t*_1_ and *t*_2_, and mutations in cluster C occurring between *t*_2_ and *t*_3_. We see the tumor composition change from consisting primarily of clone 4 (containing mutation clusters A and D) at *t*_1_, to consisting primarily of clone 3 (containing mutation clusters A, B and C) at *t*_5_.

**Figure 6:**
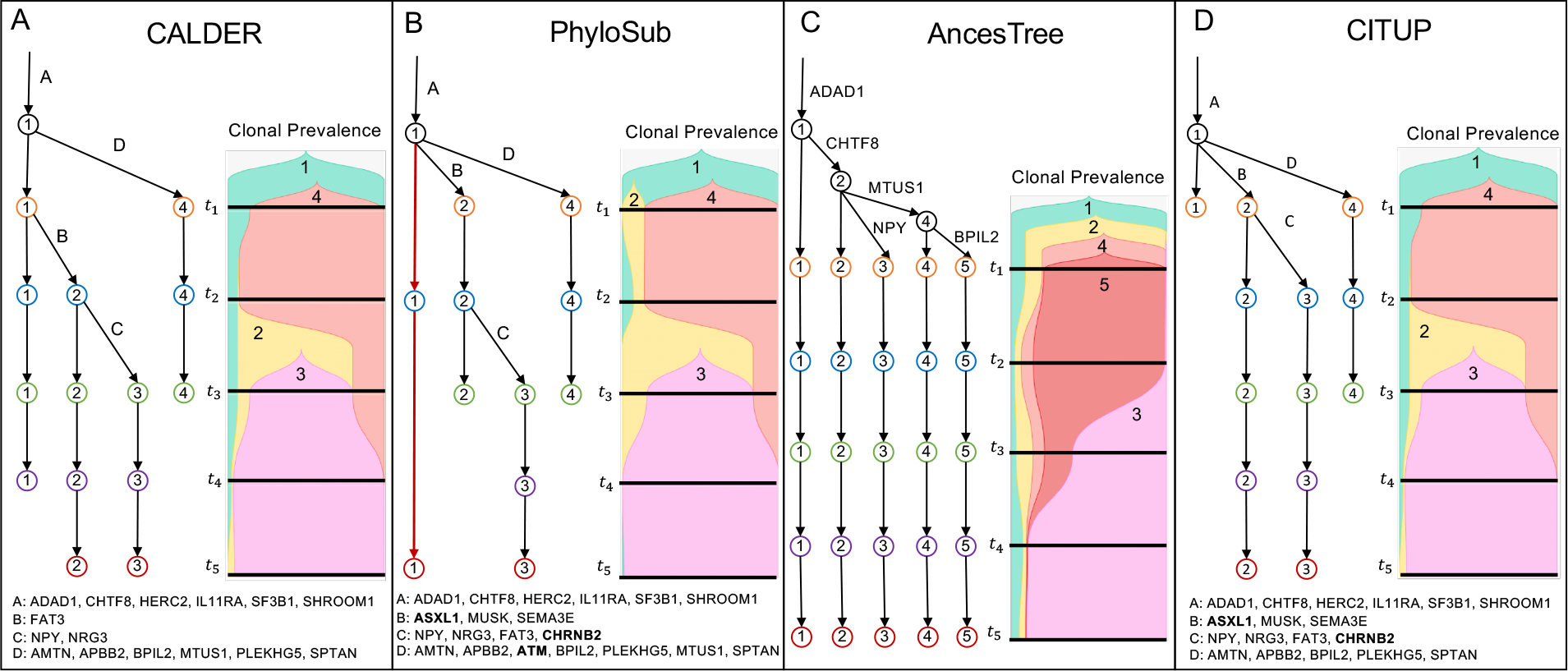
Results from (A) CALDER and previously published results from (B) PhyloSub [13], (C) AncesTree [3], and (D) CITUP [14], on CLL patient CLL003 from Schuh et al. [23]. For CALDER, PhyloSub, and CITUP, edges are labeled by mutation clusters. Mutations that CALDER and AncesTree excluded from tree construction are marked in boldface.

We compare the CALDER solution to the published results from PhyloSub [13], AncesTree [3], and CITUP [14] on this patient (Figure 6B-D). The maximum-likelihood tree inferred by PhyloSub (Figure 6B) violates longitudinal constraints (highlighted in red on the tree), with clone 1 being present only at *t*_2_ and *t*_5_.

The tree reported by AncesTree (Figure 6B) includes 5/17 unclustered mutations. The corresponding clone tree is trivially longitudinal, with all clones present at all times, and consequently implies that the entirety of the tumor’s evolution transpired before the first sample was taken. Many entries in the corresponding usage matrix are very small (highlighted in red), and the presence of these clones is not well-supported by the data. For example, the presence of clone 3 at time-point *t*_1_ is determined by the presence of mutation NPY. However, only 0.09% of the total reads (42 of 45,586) at this locus support this mutation, which is within the range of the estimated per-base sequencing error rate for Illumina sequencers of ≥ 0.1% [8]. Overall, we find that the CALDER solution for this patient is more plausible than the PhyloSub and AncesTree solutions. Interestingly, the tree inferred by CITUP (Figure 6C) is longitudinally observed (but not trivially longitudinal), despite the fact that CITUP does not explicitly enforce longitudinal constraints. It is possible that the combinatorial optimization algorithm used in CITUP favors solutions where *U* is sparse, leading to a longitudinal solution in this case.

## 4 Discussion

Longitudinal sampling is becoming more common in cancer studies, particularly in leukemias and lymphomas, where biopsies are straightforward. Even for solid tumors, ctDNA and circulating tumor cell sequencing technologies are providing the ability to monitor cancer patients over time. Longitudinal sequencing enables researchers to gain further insight into tumor evolution by revealing shifts in clonal composition over time and/or in response to treatment. Here we demonstrated how longitudinal observations inform the reconstruction of phylogenetic trees from mixed samples. We introduced CALDER, an algorithm that leverages constraints from longitudinal sampling to reduce ambiguities in the reconstruction of phylogenies from bulk DNA sequencing data. We also introduce an absence-aware clustering approach that distinguishes between mutations that are absent in a sample and those that are present in low frequency. This absence-aware clustering is particularly important for enforcing longitudinal constraints, which rely on the presence and absence of clones. In addition, absence-aware clustering is helpful when analyzing cellular migration patterns using tumor samples from distinct anatomical sites, e.g., primary tumor and metastases samples from the same patient as studied in [5]. We showed on simulated and real data that CALDER yields more plausible phylogenetic trees than existing approaches.

There are several limitations of the present approach and directions for future work. First, we focused on SNVs in diploid regions, ignoring copy number aberrations which complicate tumor phylogeny reconstruction [2, 4, 12]. While copy number aberrations are less of an issue for the leukemias studied here, it is important to extend the longitudinal constraints to include copy number aberrations. Second, as with any tumor phylogeny construction algorithm, CALDER relies on clustering of mutations into clones. While we presently cluster in advance of phylogeny reconstruction, simultaneous absence-aware clustering and phylogenetic reconstruction with CALDER is an important future goal, as has been shown previously for methods that do not use longitudinal constraints [2, 12, 13, 22]. Finally, while CALDER was developed to analyze bulk sequencing data under the infinite sites assumption, we note that the longitudinal model we have presented is general, and can be extended to analyze other data types including single-cell sequencing, methylation, and gene expression data. Some of these data types may require extensive further work to develop an algorithm that applies longitudinal constraints. However, even without this work, one can apply longitudinal constraints post-hoc to solutions of exiting algorithms to assess their plausibility. Finally, while we presented the longitudinal model in the context of cancer evolution, we note that no part of the model or algorithm is specific to cancer, and CALDER can be applied to other evolving systems. One promising application of CALDER is to study the shifting composition of bacterial populations from longitudinal metagenomic samples [6].

## Acknowledgments

This work is supported by a US National Institutes of Health (NIH) grants R01HG007069 and U24CA211000 and US National Science Foundation (NSF) CAREER Award (CCF-1053753) to BJR.

## References

[1] HX Dang, BS White, SM Foltz, CA Miller, J Luo, RC Fields, and CA Maher. Clonevol: clonal ordering and visualization in cancer sequencing. Annals of Oncology, 28(12):3076–3082, 2017.

[2] Amit G Deshwar, Shankar Vembu, Christina K Yung, Gun Ho Jang, Lincoln Stein, and Quaid Morris. Phylowgs: reconstructing subclonal composition and evolution from whole-genome sequencing of tumors. Genome biology, 16(1):35, 2015.

[3] Mohammed El-Kebir, Layla Oesper, Hannah Acheson-Field, and Benjamin J Raphael. Reconstruction of clonal trees and tumor composition from multi-sample sequencing data. Bioinformatics, 31(12):i62–i70, 2015.

[4] Mohammed El-Kebir, Gryte Satas, Layla Oesper, and Benjamin J Raphael. Inferring the mutational history of a tumor using multi-state perfect phylogeny mixtures. Cell systems, 3(1):43–53, 2016.

[5] Mohammed El-Kebir, Gryte Satas, and Benjamin J Raphael. Inferring parsimonious migration histories for metastatic cancers. cancer, 2:5, 2018.

[6] Karoline Faust, Leo Lahti, Didier Gonze, Willem M de Vos, and Jeroen Raes. Metagenomics meets time series analysis: unraveling microbial community dynamics. Current opinion in microbiology, 25:56–66, 2015.

[7] Harold N Gabow and Eugene W Myers. Finding all spanning trees of directed and undirected graphs. Society for Industrial and Applied Mathematics, 7(3):280–287, 1978.

[8] Travis C Glenn. Field guide to next-generation dna sequencers. Molecular ecology resources, 11(5):759–769, 2011.

[9] Malachi Griffith, Christopher A Miller, Obi L Griffith, Kilannin Krysiak, Zachary L Skidmore, Avinash Ramu, Jason R Walker, Ha X Dang, Lee Trani, David E Larson, et al. Optimizing cancer genome sequencing and analysis. Cell systems, 1(3):210–223, 2015.

[10] Dan Gusfield. Haplotyping as perfect phylogeny: conceptual framework and efficient solutions. Proceedings of the sixth annual international conference on Computational biology, pages 166–175, 2002.

[11] Daniel A Haber and Victor E Velculescu. Blood-based analyses of cancer: circulating tumor cells and circulating tumor dna. Cancer discovery, 4(6):650–661, 2014.

[12] Yuchao Jiang, Yu Qiu, Andy J Minn, and Nancy R Zhang. Assessing intratumor heterogeneity and tracking longitudinal and spatial clonal evolutionary history by next-generation sequencing. Proceedings of the National Academy of Sciences, 113(37):E5528–E5537, 2016.

[13] Wei Jiao, Shankar Vembu, Amit G Deshwar, Lincoln Stein, and Quaid Morris. Inferring clonal evolution of tumors from single nucleotide somatic mutations. BMC bioinformatics, 15(1):35, 2014.

[14] Salem Malikic, Andrew W McPherson, Nilgun Donmez, and Cenk S Sahinalp. Clonality inference in multiple tumor samples using phylogeny. Bioinformatics, 31(9):1349–1356, 2015.

[15] Christopher A Miller, Brian S White, Nathan D Dees, Malachi Griffith, John S Welch, Obi L Griffith, Ravi Vij, Michael H Tomasson, Timothy A Graubert, Matthew J Walter, et al. Sciclone: inferring clonal architecture and tracking the spatial and temporal patterns of tumor evolution. PLoS computational biology, 10(8):e1003665, 2014.

[16] Ferran Nadeu, Julio Delgado, Cristina Royo, Tycho Baumann, Tatjana Stankovic, Magda Pinyol, Pedro Jares, Alba Navarro, David Martín-García, Sílvia Beà, et al. Clinical impact of clonal and subclonal tp53, sf3b1, birc3, notch1 and atm mutations in chronic lymphocytic leukemia. Blood, pages blood–2015, 2016.

[17] Peter C Nowell. The clonal evolution of tumor cell populations. Science, 194(4260):23–28, 1976.

[18] Victoria Popic, Raheleh Salari, Iman Hajirasouliha, Dorna Kashef-Haghighi, Robert B West, and Serafim Batzoglou. Fast and scalable inference of multi-sample cancer lineages. Genome biology, 16(1):91, 2015.

[19] Johannes G Reiter, Alvin P Makohon-Moore, Jeffrey M Gerold, Ivana Bozic, Krishnendu Chatterjee, Christine A Iacobuzio-Donahue, Bert Vogelstein, and Martin A Nowak. Reconstructing metastatic seeding patterns of human cancers. Nature Communications, 8:14114, 2017.

[20] Matthew JJ Rose-Zerilli, Jane Gibson, Jun Wang, William Tapper, Zadie Davis, Helen Parker, Marta Larrayoz, Helen McCarthy, Renata Walewska, Jade Forster, et al. Longitudinal copy number, whole exome and targeted deep sequencing of‘good risk’ighv-mutated cll patients with progressive disease. Leukemia, 30(6):1301, 2016.

[21] Andrew Roth, Jaswinder Khattra, Damian Yap, Adrian Wan, Emma Laks, Justina Biele, Gavin Ha, Samuel Aparicio, Alexandre Bouchard-Côté, and Sohrab P Shah. Pyclone: statistical inference of clonal population structure in cancer. Nature methods, 11(4):396, 2014.

[22] Gryte Satas and Benjamin J Raphael. Tumor phylogeny inference using tree-constrained importance sampling. Bioinformatics, 33(14):i152–i160, 2017.

[23] Anna Schuh, Jennifer Becq, Sean Humphray, Adrian Alexa, Adam Burns, Ruth Clifford, Stephan M. Feller, Russell Grocock, Shirley Henderson, Irina Khrebtukova, Zoya Kingsbury, Shujun Luo, David McBride, Lisa Murray, Toshi Menju, Adele Timbs, Mark Ross, Jenny Taylor, and David Bentley. Monitoring chronic lymphocytic leukemia progression by whole genome sequencing reveals heterogeneous clonal evolution patterns. Blood, 120(20):4191–4196, November 2012.

[24] Francesco Strino, Fabio Parisi, Mariann Micsinai, and Yuval Kluger. Trap: a tree approach for finger-printing subclonal tumor composition. Nucleic acids research, 41(17):e165–e165, 2013.

[25] Robert Tibshrani. Regression shrinkage and selection via the lasso. Journal of the Royal Statistical Society, 58(1):267–288, 1996.

[26] Habil Zare, Junfeng Wang, Alex Hu, Kris Weber, Josh Smith, Debbie Nickerson, ChaoZhong Song, Daniela Witten, C Anthony Blau, and William Stafford Noble. Inferring clonal composition from multiple sections of a breast cancer. PLoS computational biology, 10(7):e1003703, 2014.

